# Accounting for measurement error to assess the effect of air pollution on omic signals

**DOI:** 10.1101/504001

**Authors:** Erica Ponzi, Paolo Vineis, Kian Fan Chung, Marta Blangiardo

## Abstract

Studies on the effects of air pollution and more generally environmental exposures on health require measurements of pollutants, which are affected by measurement error. This is a cause of bias in the estimation of parameters relevant to the study and can lead to inaccurate conclusions when evaluating associations among pollutants, disease risk and biomarkers. Although the presence of measurement error in such studies has been recognized as a potential problem, it is rarely considered in applications and practical solutions are still lacking.

In this work, we formulate Bayesian measurement error models and apply them to study the link between air pollution and omic signals. The data we use stem from the “Oxford Street II Study”, a randomized crossover trial in which 60 volunteers walked for two hours in a traffic-free area (Hyde Park) and in a busy shopping street (Oxford Street) of London. Metabolomic measurements were made in each individual as well as air pollution measurements, in order to investigate the association between short-term exposure to traffic related air pollution and perturbation of metabolic pathways. We implemented error-corrected models in a classical framework and used the flexibility of Bayesian hierarchical models to account for dependencies among omic signals, as well as among different pollutants. Models were implemented using traditional MCMC simulative methods as well as integrated Laplace approximation.

The inclusion of a classical measurement error term resulted in variable estimates of the association between omic signals and traffic related air pollution (TRAP) measurements, where the direction of the bias was not predictable a priori. The models were successful in including and accounting for different correlation structures, both among omic signals and among different pollutant exposures. In general, more associations were identified when the correlation among omics and among pollutants were modeled, and their number increased when a measurement error term additionally included in the multivariate models (particularly for the associations between metabolomics and NO_2_).

## Introduction

Health effects of air pollution are a major public health issue and have received increasing attention over the past decades (Elliott et al. 1992; Pekkanen and Pearce 2001; Baker and Nieuwenhuijsen 2008). In this context, the reliable estimation of associations between environmental exposures and health conditions requires the collection of a huge amount of exposure data, which is often impractical and subject to several sources of error or imprecision. This can lead not only to bias in the estimation of parameters relevant to the study but also to inaccurate conclusions when evaluating associations among pollutants, disease risk and biomarkers. Although the presence of measurement error in such studies has been discussed in the recent literature and is now recognized as a potentially serious problem (Rhomberg et al. 2011; Edwards and Keil 2017), it is often not accounted for in standard analyses.

Different approaches to the problem have been adopted and different methods and techniques are available in the literature, for instance Mallick, Hoffman, and Carroll (2002) suggested a semi-parametric approach to different types of error in radiation data, Gryparis, Coull, and Schwartz (2007) proposed a Bayesian hierarchical model to retrieve error-free estimates of the health effect, Baxter et al. (2010) a quantification of measurement error effect via validation studies, and Goldman et al. (2011) proposed a method to account for the error in time-series studies on air pollution. Studies focusing on the effect of traffic related air pollution (TRAP) are particularly challenging and often rely on surrogate measures of pollutants, as well as on the approximation of personal exposures. Zeger et al. (2000) identified three main sources of error in the TRAP exposure assessment, namely (a) the difference between measured and true ambient exposure levels, (b) the difference between aggregate personal exposure and the exposure of a given individual, mostly due to approximation and classified as Berkson error, and (c) the difference between ambient concentration and average personal exposure, classified as a classical error. In this context, different methods have been proposed and a wide set of measurement error techniques have been employed, including among others the simulation extrapolation algorithm (Alexeeff, Carroll, and Coull 2016), regression calibration and validation data (Van Roosbroeck et al. 2006) and Bayesian methods (Gryparis, Coull, and Schwarzt 2007; Gryparis et al. 2009). On the other hand, none of these studies was applied to omic data, as they focused on estimating disease risks and did not include any molecular data.

In the present study, we propose to apply techniques to correct for measurement error in environmental exposures when considering their association with high-throughput molecular data. This is particularly challenging due to the high dimensionality of the data, as well as to the correlation among omics sampled from the same individual. We use a Bayesian framework to address the problem, which provides a very flexible way to account for measurement error and to model different error types and dependency structures in the data. In particular, Bayesian hierarchical models seem ideal in these contexts, as they provide a straightforward way to include dependency between exposures, but also between different response variables.

Moreover, the possibility to include prior knowledge on the error components can result in better models and more accurate estimations and the possibility to model several fixed and random effects, as well as different link functions, adds flexibility and general applicability to the methods.

In this paper, we apply this approach to the Oxford Street II study, a randomized crossover trial where omics and air pollution measurements were employed to investigate the association between short-term exposure to TRAP and perturbation of different omic signals (Sinharay et al. 2017; van Veldhoven et al. 2019). We implemented error-corrected models in a classical measurement error framework and generalized such models to account for dependencies among pollutants, as well as among response omic variables. This provides a novel way of dealing with high-dimension omic data, by including them into a Bayesian hierarchical formulation. The possibility to model more omic signals at the same time also allows to account for dependency among signals. Moreover, the inclusion of a measurement error term, which is straightforward and flexible thanks to the hierarchical formulation, has not been proposed so far in the presence of high-throughput biological data.

We implement our models using `MCMC` in `JAGS`, but to increase the speed of the computation, we also use the integrated nested Laplace approximation approach (INLA) (Rue, Martino, and Chopin 2009).

The remainder of this paper is structured as follows: we first describe the study and the model to assess the association between different air pollutants and omic measurements, focusing on metabolic pathways. The paper then illustrates the Bayesian hierarchical model we formulate to account for measurement error by including a classical error. We expand such model to a multi-response model, accounting for a dependency structure among different omic signals, and to a multi-variate model to account for dependency among different pollutants. We then show the results based on the data set from the Oxford Street II study and finally conclude with several discussion points and potential expansion of the proposed method.

## Methods

### Metabolic pathways in the Oxford Street II Study

#### The study

The data we use here stem from the Oxford Street II Study, a randomized crossover trial within the EXPOsOMICS consortium (Vineis et al. 2017). In this study, 60 volunteers walked for two hours in a traffic-free area (Hyde Park) and in a busy shopping street (Oxford Street) of London. The walking experiments were performed on non-rainy weekdays only, from November to March, to avoid confounding from rain or pollen. Participants were divided into three groups: 1) healthy volunteers (n=20) with a normal lung function and without a history of ischemic heart disease (IHD); 2) patients with chronic obstructive pulmonary disease (COPD) (n=20), without a history of IHD; and 3) patients with clinically stable IHD over the past six months (n=20) without COPD. All current smokers or former smoker for less than 12 months were excluded, as well as people with high occupational levels of TRAP. Information on age, sex, BMI, blood pressure, distance walked, diet and medication use was collected for each participant.

For each individual and each exposure session, three blood samples were collected: two hours before walking, two hours after walking and 24 hours after walking; at the same time TRAP measurements were taken, namely on black carbon (CBLK), nitrogen dioxide (NO_2_), particulate matter PM_10_ and PM_25_. Such measurements are likely to suffer from classical measurement error, as they were collected using a portable size-selective airborne particle sampler and therefore rely on each patient’s precision and accuracy, besides being potentially subject to the sources of error in TRAP identified by Zeger et al. (2000) and reported above. Moreover, as reported in Sinharay et al. (2017), “NO_2_ concentrations were taken from a stationary monitoring site on Oxford Street repeatedly passed during walks on Oxford Street. Because no monitoring was available in Hyde Park, NO_2_ concentrations were taken from the nearest representative location sited in a school playground”. Finally, real-time measurements of noise, temperature and relative humidity were obtained at each exposure session. On such samples, different untargeted omic analyses were performed, including metabolomic analyses, on which we focus here. A more detailed description of the study is given in Sinharay et al. (2017), where it is also reported that the study was approved by the UK National Research Ethics Service and that informed written consent was obtained from all participants. Van Veldhoven et al. (2019) conducted analyses to assess the short term metabolomic changes due to exposure to TRAP and identified various associations between air pollution concentrations and levels of metabolites in blood.

#### The model

The association between metabolite levels and TRAP exposures was assessed in a mixed model framework, using a Bayesian approach and including random effects for the individual, as well as for the location and time point of each measurements. Fixed effects were sex, age, body mass index (BMI) and health group, as well as annual and instantaneous measurements of the exposure of interest. The four exposures reported above were considered separately.

The model was formulated as follows:

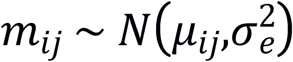

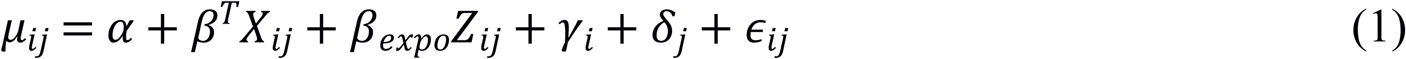

with *m*_*ij*_ being the metabolic feature of individual *i* at measurement *j, Z* the exposure, **X** including sex, age, BMI and health group, as well as the annual measurement of the exposure of interest and 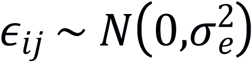. Random effects were included for the individual *γ* and for the interaction *δ* between time point and location, which ranged from 1 to 6 (three time points per location) for each individual. Normally distributed priors with mean equal to zero and variance equal to 1000 were assigned to the regression coefficients **β**, and vague inverse gamma priors with shape and scale equal to 0.01 were assigned to the precisions of the random effects. A graphical representation of the model is reported in Figure 1, while a similar, frequentist approach to the same model was used in van Veldhoven et al. (2019). Given the extreme high dimensionality of the omics dataset we pre-selected only the signals which reported p-values smaller than 0.5 in the frequentist analysis.

**Figure1:**
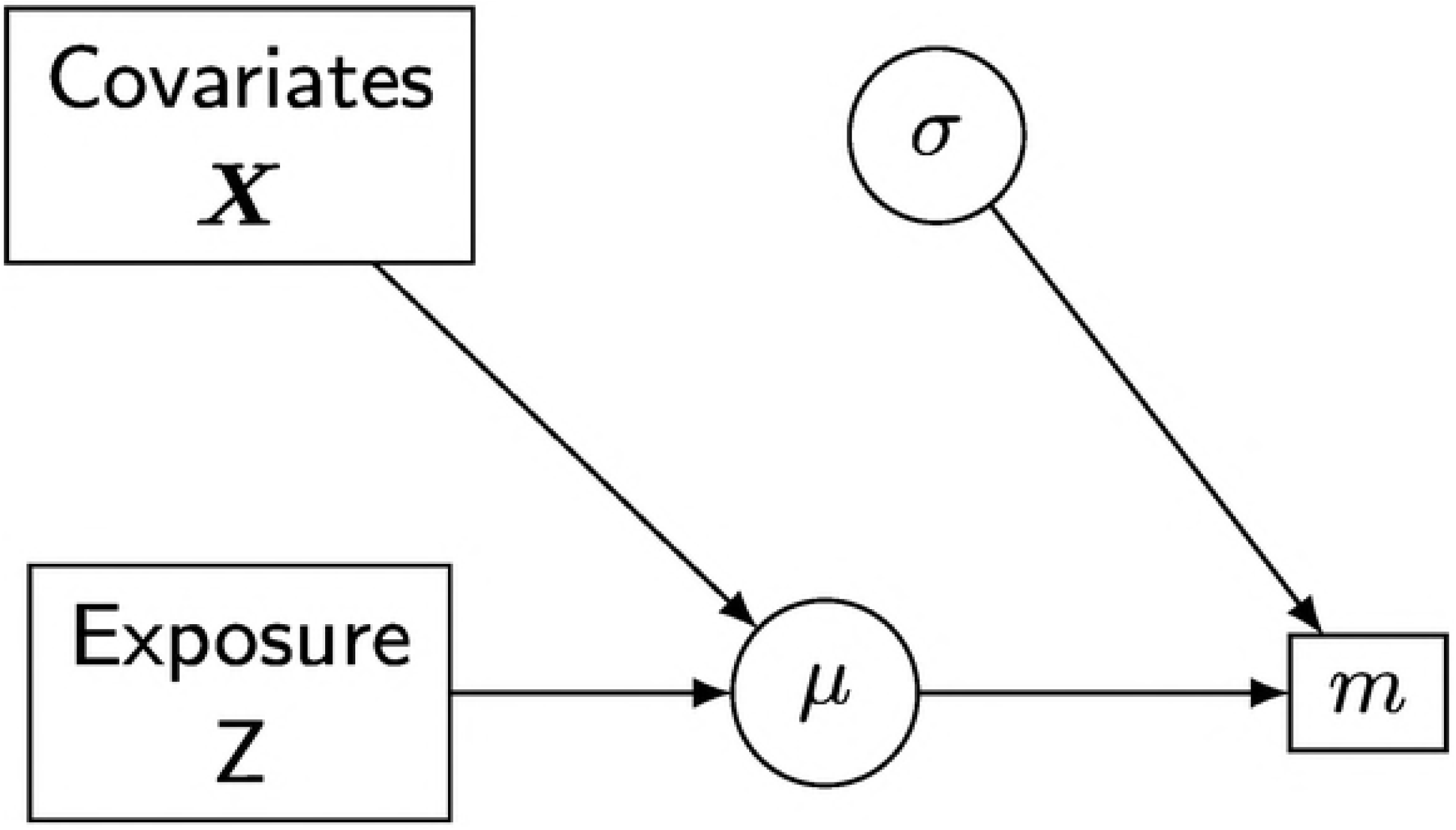
Bayesian formulation of the basic model to assess association between metabolic levels and TRAP exposure. Squares are used to indicate data, while circles represent random variables or deterministic relationships.

To determine whether there was evidence of an association between the signal and the exposure of interest, we used posterior predictive probabilities to calculate Bayesian p-values and corrected for multiple testing using the Bayesian False Discovery Rate (FDR) (Newton et al. 2004; Ventrucci, Scott, and Cocchi 2011).

This model, and all models reported in this paper, were formulated using `JAGS` and `INLA` and the code to implement them is reported in the supplementary material. From now on, we will denote this model as the “naive” model, to indicate the fact that it does not account for the presence of measurement error, while we will use “corrected” for the model which includes a component for measurement error.

### Classical measurement error

To model the presence of measurement error, we included an additional component in the model by formulating a classical measurement error for the different pollutants, i.e. assuming that the exposure variable **Z** can be observed only via a proxy **W**, such that

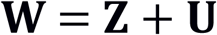

with 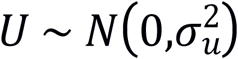. In the presence of a classical measurement error, an attenuation of the effect of the error-prone variable is expected, as the presence of the additional error variance biases the estimates of the regression parameters towards zero (Carroll et al. 2006). In fact, bearing in mind that 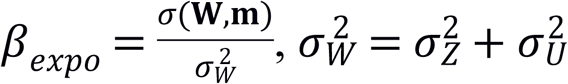, and assuming that the error in **W** is independent of **m** and of any other variables, it is quite straightforward to see that the error-prone regression parameter will be given by

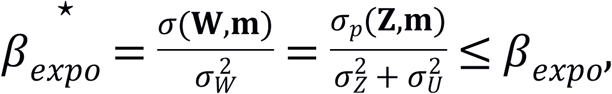

and that the quantity that is estimated is 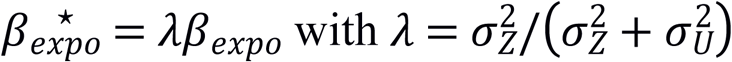. It is important to underline that, even if an attenuation is usually expected, upward bias is also a possible consequence of classical measurement error even in relatively simple models, due for example to a correlation between covariates (Carroll et al. 2006). It is therefore necessary to disentangle the error variance from the true variance measured by the proxy in order to obtain unbiased estimates of the regression coefficients. To do so, we formulated a *Bayesian hierarchical model*, as the hierarchical formulation and the possibility to include prior knowledge provide a flexible way to model measurement error (Stephens and Dellaportas 1992; Richardson and Gilks 1993).

Such formulation was included in the model by adding a latent variable for the exposure, namely a normally distributed variable with mean equal to 0 and variance equal to the error variance. The underlying level of the structure was instead the same as in model (1), resulting in the following hierarchical structure:

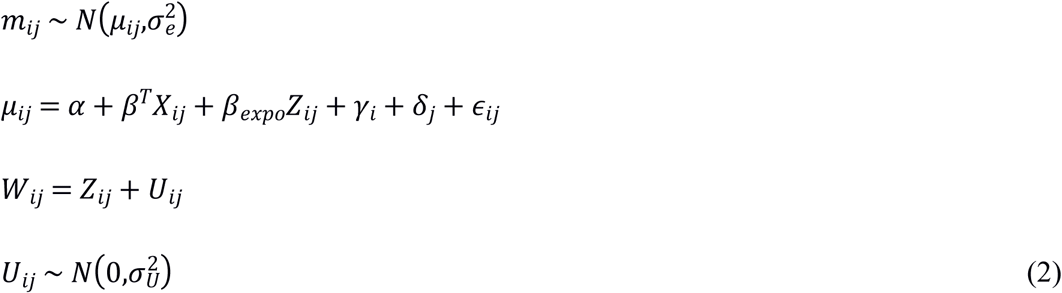

A vague gamma prior was used for the error variance, with shape and scale parameters equal to 0.01. For all the other variance components and regression coefficients, the same priors were used as in the naive model reported in the previous section.

A graphical representation of the model is reported in Figure 2.

**Figure2:**
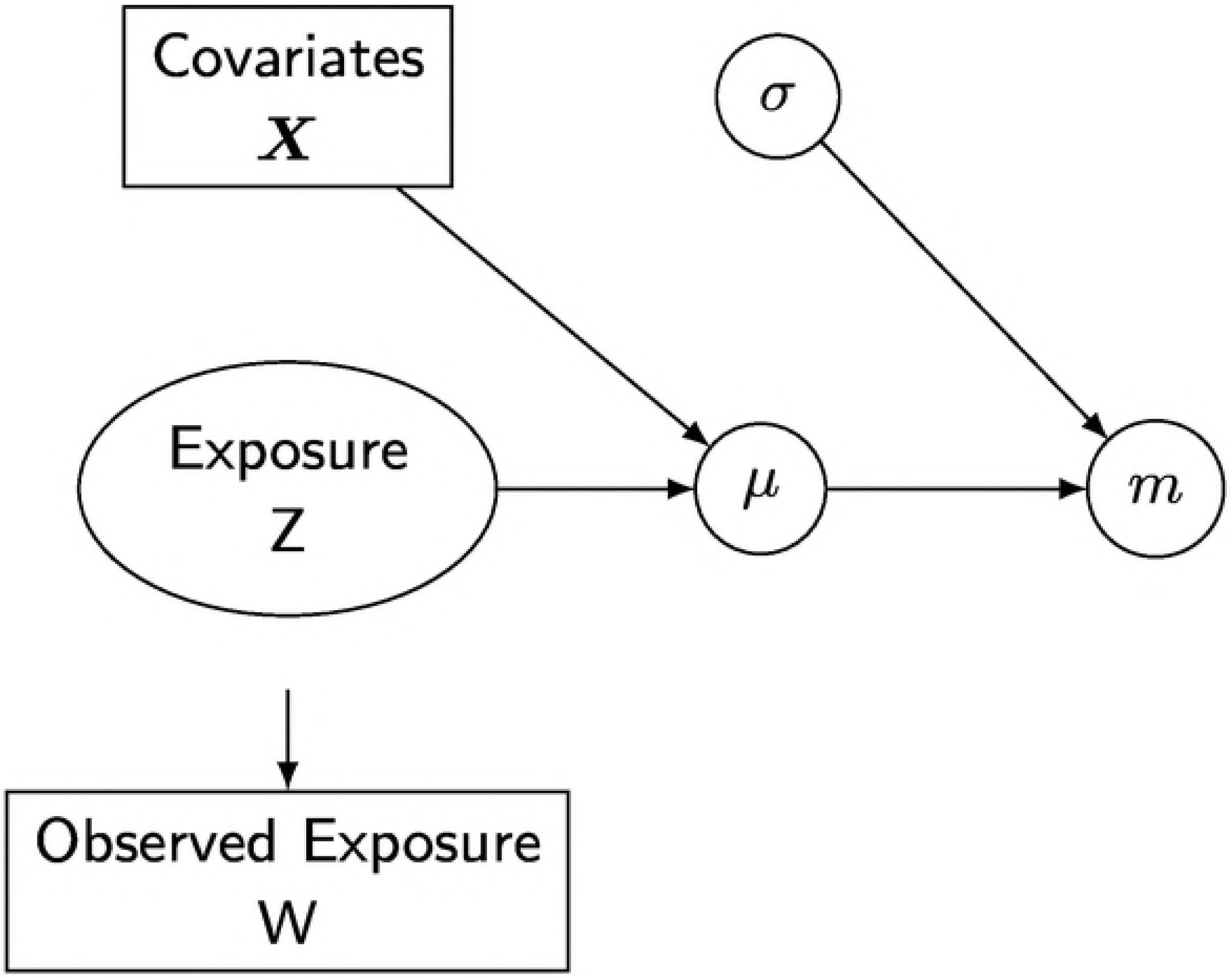
Bayesian formulation of the classical error model. Z is not observed, W is the observed proxy for the exposure

## Multivariate models

### Dependency among omic signals

Another source of imprecision and possible bias in the assessment of the association between omic signals and TRAP exposure is potentially given by the formulation of independent models for each omic feature. Dependency across metabolic features is very likely to occur in practice, first of all because 5749 different features are sampled and analysed from the same 60 individuals, and second because they all reflect metabolic pathways and phenomena that are highly correlated in each individual. The flexibility and the layer structure of Bayesian hierarchical models makes it straightforward to account for such dependency, namely by using a multivariate response for the omic signals and introducing dependency in their variance covariance structure. The resulting model is a generalization of model (2) and was formulated as follows:

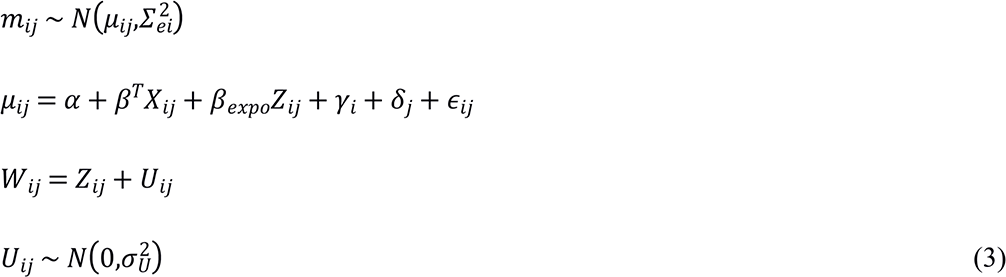

where the response variable followed a multivariate normal distribution **m** ∼ **N**(0,**∑**_e_), with **∑**_e_ denoting the covariance matrix of the omics signals and all variances were given a gamma prior with shape and scale equal to 0.01, in consistency with the univariate model. Note that this formulation with multivariate signals does not require any correction for multiple testing, unlike the initial univariate model, as it naturally accounts for dependency across omics. Therefore, no Bayesian FDR was used in this context and associations were simply assessed by means of posterior predictive distributions.

To model the presence of the error in the exposure variable, we used the same reasoning as before and introduced a latent variable for the exposure, namely a normally distributed variable with mean equal to 0 and variance equal to the error variance. The error structure was again given by a classical error model and a vague gamma prior was used for the error variance, with shape and scale parameters equal to 0.01, again in consistency with the formulation and priors used in the univariate model.

### Dependency among different exposures

Dependency structures are very likely to occur in such studies also between different environmental exposures, as different works in the literature recently pointed out (Coker et al. 2016; Huang and Lee 2018). In particular, all the TRAP measurements show a general common trend based on traffic conditions, as well as weather, humidity and temperature, but also based on other confounding factors registered in each particular day. Moreover, individual-specific factor can cause a variation in all pollutants at the same time, affecting in our specific case the measurements taken before and after the walk experiment. Such dependency appears to be very frequent between particulate matters of different sizes, as they are induced by similar mechanisms and conditions, besides being measured using the same techniques and instruments, and between PMs and black carbon. The Bayesian hierarchical structures employed in the previous sections can be further generalized to account for such correlation, by modelling more TRAP exposures at the same time. The resulting model was obtained as a generalization of (3) and formulated as follows:

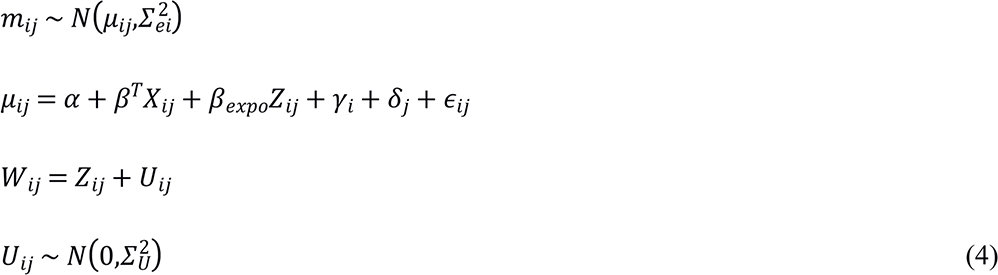

All coefficients and variables were modeled as previously described and the same priors were given to all of them. Again, the flexible formulation of Bayesian hierarchical structures allows to include as many pollutants as desired and to inform the model on their correlation and variances based on our knowledge. In analogy with the first univariate model, we added a latent variable for the multivariate exposure, following a normal distribution with mean equal to 0 and variance equal to the error variance. To account for dependency among pollutants, *U* was multivariate and modeled as a vector with four dimensions, corresponding to the four different pollutants. The variance covariance matrix of the multivariate exposure, as well as of the multivariate error, was given an uninformative Wishart prior to account for correlation among pollutants.

## Results

As expected, the inclusion of a classical measurement error term resulted in different estimates of the association between omic signals and TRAP measurements, compared to the naive model which does not include such term. Note that the presence of classical measurement error in pollutant measures can cause bias in different directions, and that the effect, as well as the direction of the error correction, is not evident a priori. Figure 5 shows the estimates of regression parameters *β*_*expo*_ obtained by models (1) and (2) for the selected omics. The error corrected models reported stronger associations between TRAP measurements and metabolomic signals, although with higher uncertainty, which reflects the uncertainty on the error component. While the naive models identified respectively 0, 2, 5 and 24 association for CBLK, PM_25_, PM_10_, NO_2_, the error corrected models only identified 0, 1, 3 and 5 associations respectively, reflecting the propagation of uncertainty about the error into the posterior distributions of parameters of interest. These signals included a signal from the phenylalanine cluster already found to be associated with NO_2_ in van Veldhoven et al. (2019), as well as an unknown signal whose association with PM_10_ was also detected by van Veldhoven et al. (2019). This also shows that some associations might be affected by an opposite impact and that attenuation is not the only possible consequence of measurement error. This is also reflected by the width of credible intervals which naturally increases when accounting for measurement error, as a consequence of the uncertainty about this component. When more information is available about the prior distribution of the error, such uncertainty will most likely decrease and potentially smaller credible intervals will be identified.

**Figure 3:**
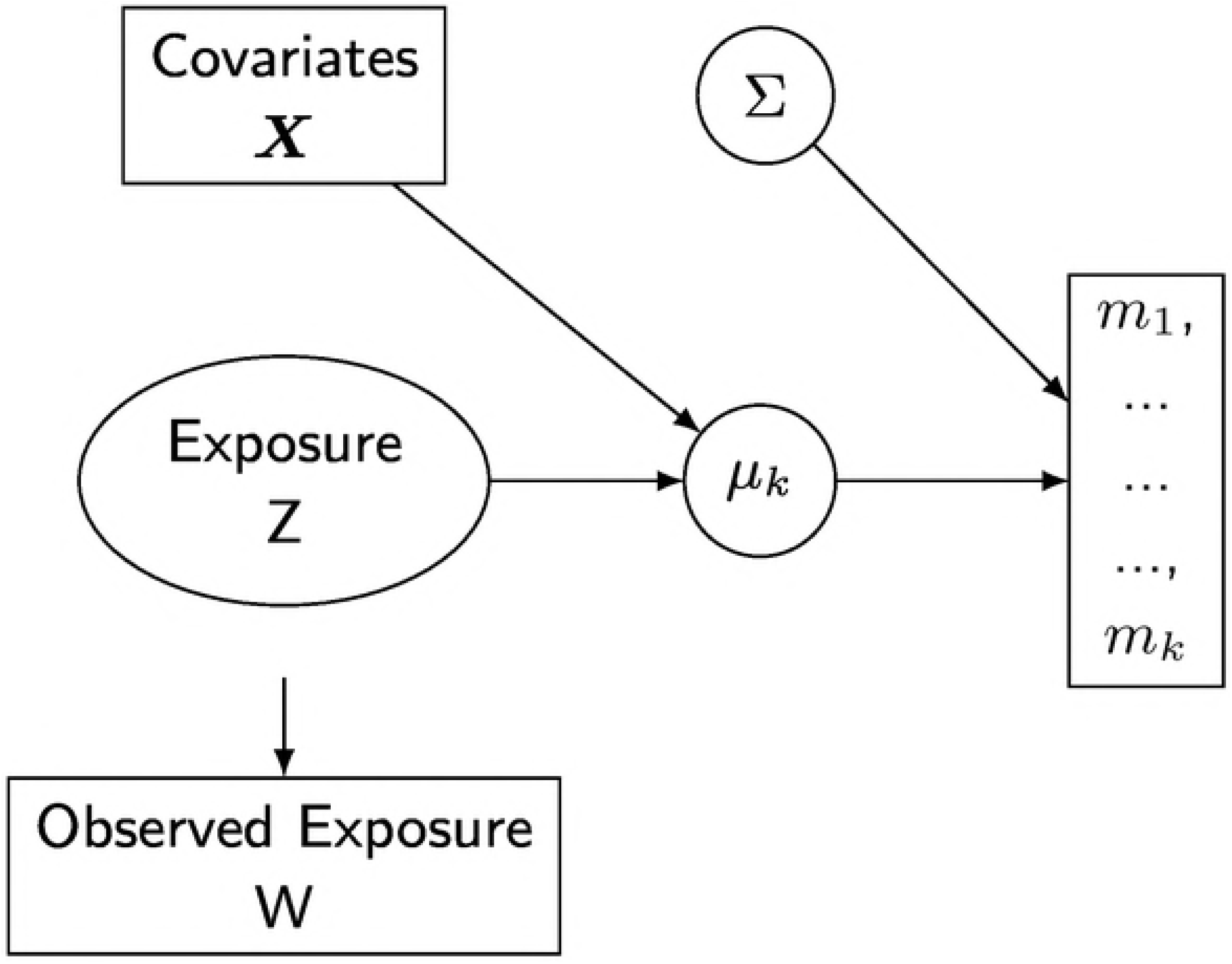
Bayesian formulation of the multivariate model with classical measurement error.

**Figure 4:**
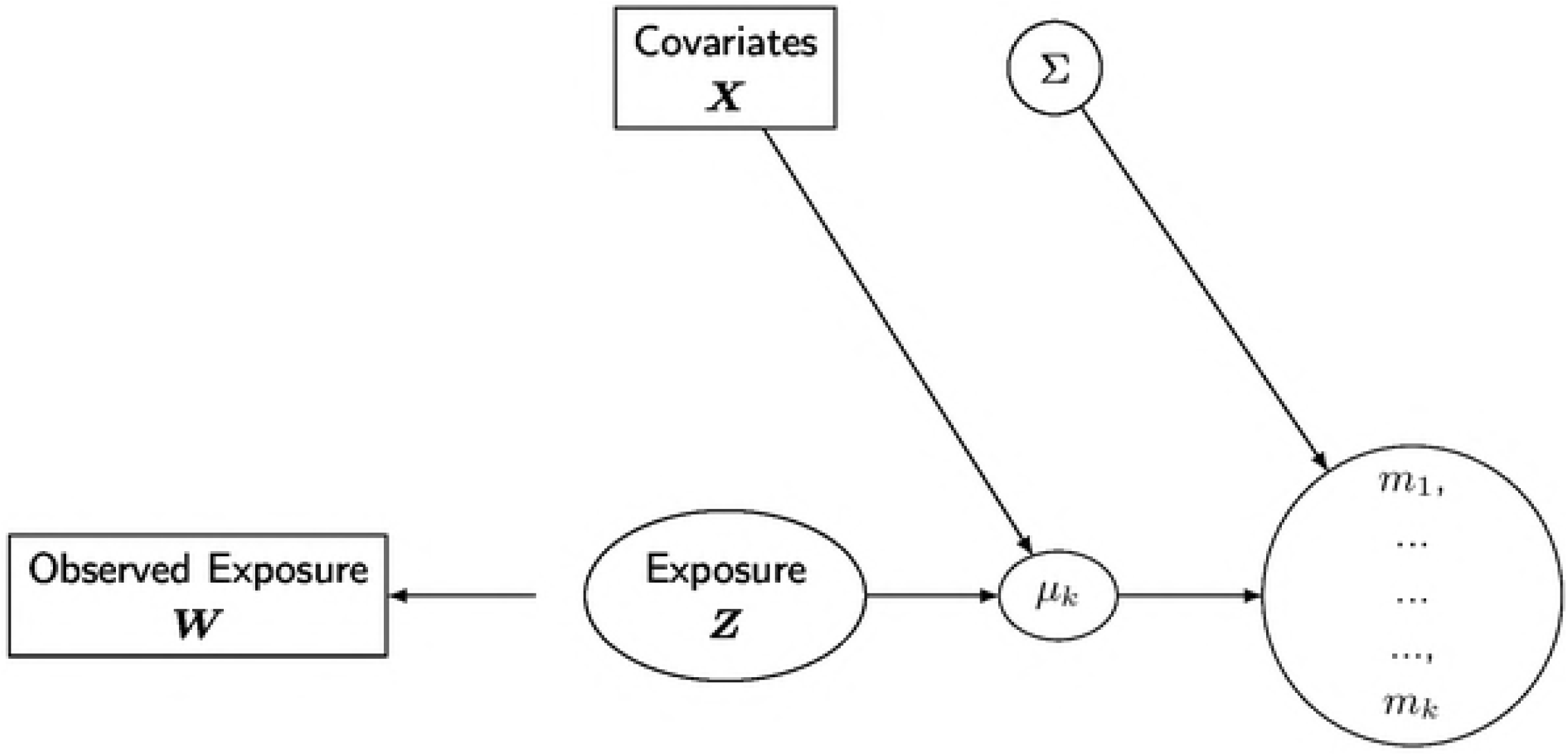
Bayesian formulation of the multivariate model with classical measurement error and multivariate exposure.

**Figure 5:**
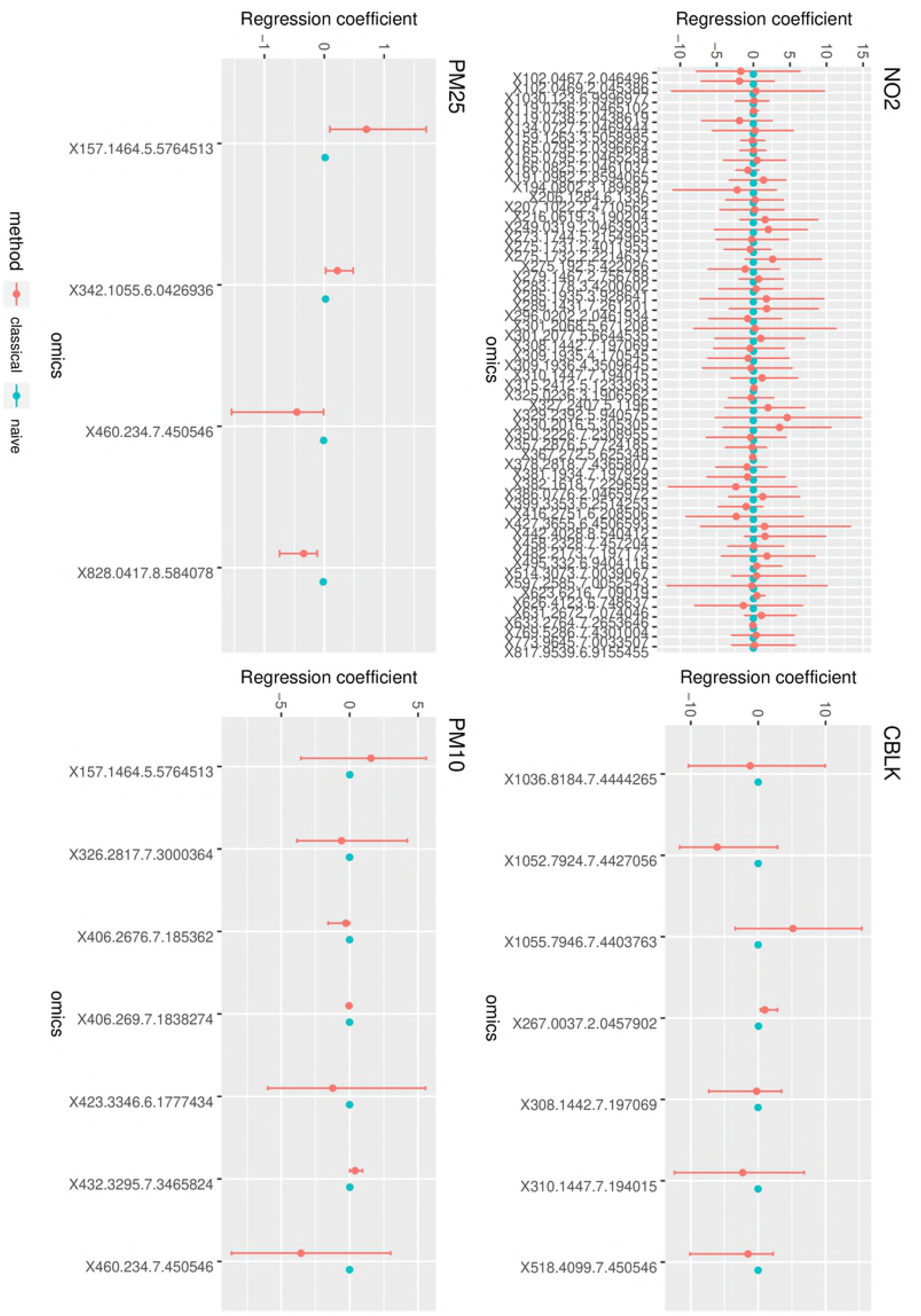
Regression coefficients with and without classical error modeling in JAGS. Estimates are reported with their 95% confidence intervals.

Such correction showed a more defined pattern when accounting for the dependency among different omic signals. In this case, the error corrected model led to estimates of *β*_*expo*_ which were constantly lower than the estimates from the naive models for most omic signals. Figure 6 shows the estimates of regression parameters *β*_*expo*_ obtained by model (3) with and without including a level for the error component. In this case the naive model identified respectively 1, 2, 2 and 4 associations for CBLK, PM_25_, PM_10_, NO_2_, while error corrected model identified respectively 6, 1, 4 and 51 associations. Note that the inclusion of a dependency structure among omic signals and of an error term in the model identifies more associations than the univariate models, because the presence of such dependencies can obscure associations among omic signals and TRAP measurements.

**Figure 6:**
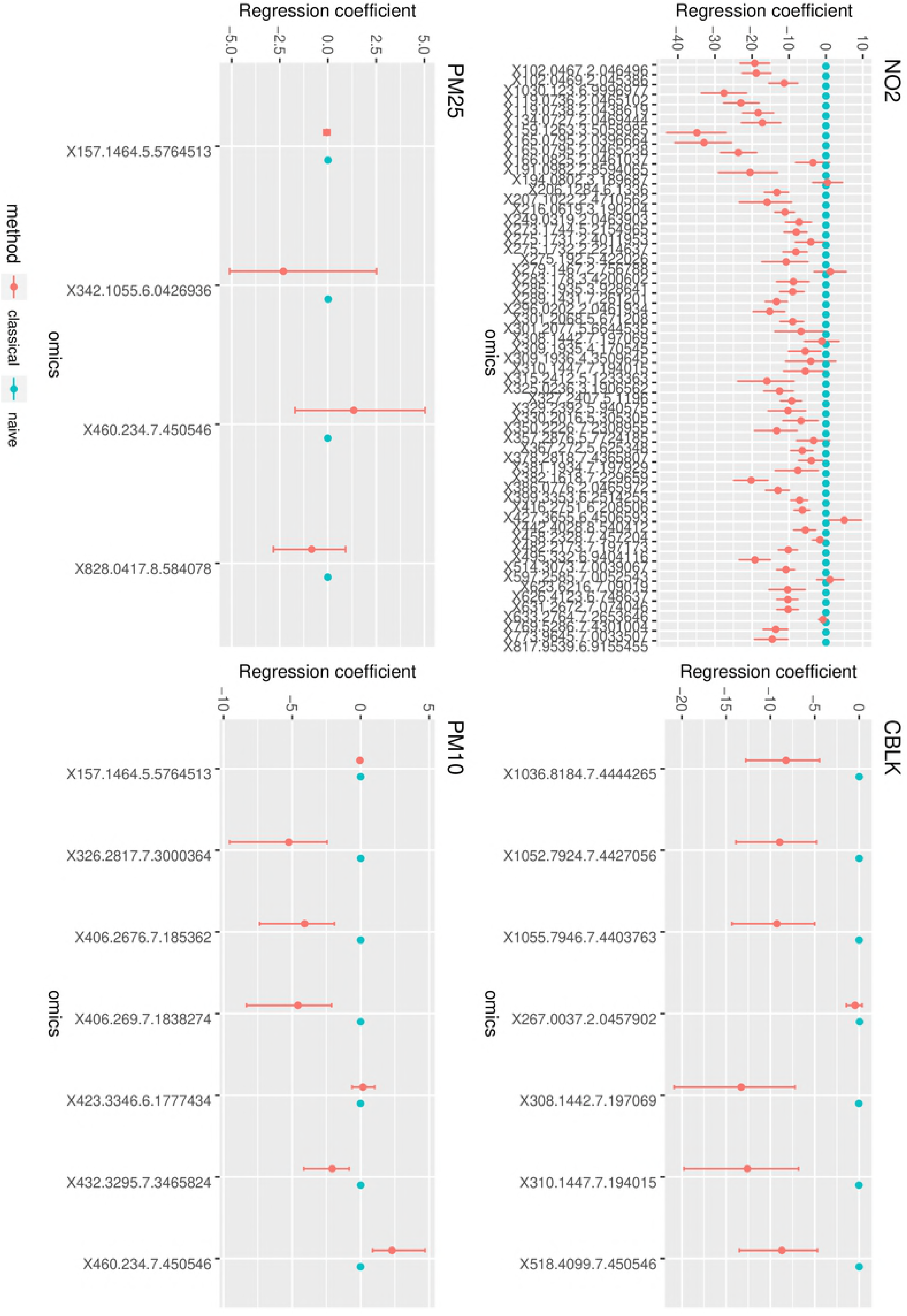
Regression coefficients with and without classical error modeling in JAGS. Estimates are reported with their confidence intervals. A correlated structure among omic signals is assumed.

When accounting for dependency among signals and among pollutants, all signals that were associated with any of the pollutants were considered. The correction introduced by the error corrected models was in most cases higher than in the previous models. Accounting for dependency among pollutants and for correlated measurement errors also identified more associations than the other models, leading to higher estimates also between signals and pollutants that were not associated in the first cases. Figure 7 shows the estimates of regression parameters *β*_*expo*_ obtained by model (4) with and without including a level for the error component. In this case the naive model identified respectively 18, 0, 25 and 16 association for CBLK, PM_25_, PM_10_, NO_2_, while error corrected model identified respectively 7, 1, 22 and 52 associations.

**Figure 7:**
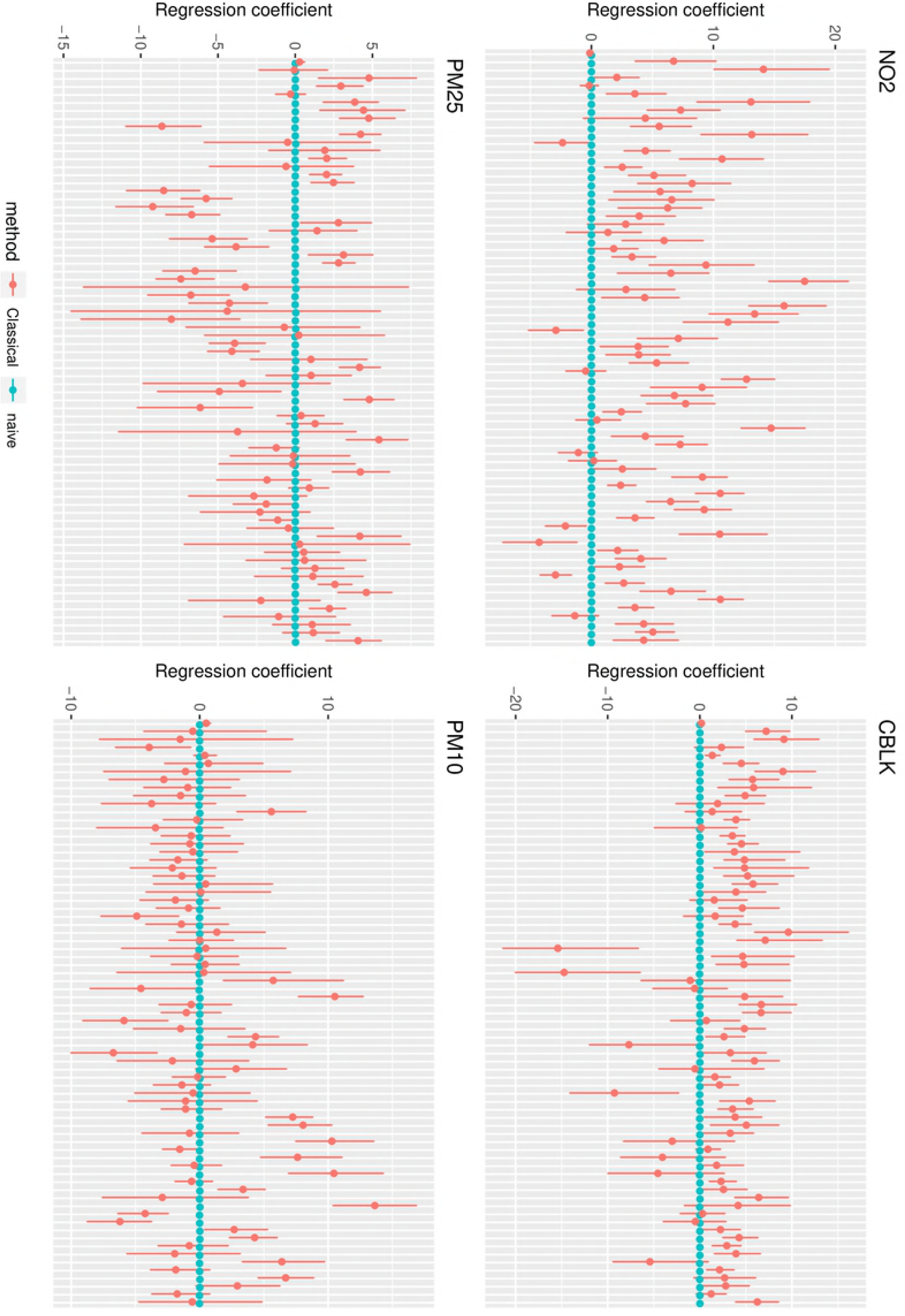
Regression coefficients with and without classical error modeling in JAGS. Estimates are reported with their confidence intervals. Dependencies among omic signals and among different TRAP exposures are modeled, as well as dependency among error components on different pollutants in the corrected model. All signals that were associated with any of the pollutants are reported for all pollutants.

This reflects the fact that ignoring the presence of measurement error can obscure associations between signals and pollutants and lead to an underestimation of effect sizes, as well as identifying potentially false associations. On the other hand, models including correlation structures identify even stronger effect sizes when corrected for measurement error, as well as a generally higher number of significant associations between signals and pollutants, which might be obscured when ignoring the presence of bias. In general, more associations were identified when the correlation among omics and among pollutants were modeled, and their number increased when a measurement error term was additionally included in the multivariate models.

## Discussion

We implemented Bayesian hierarchical models to account for the presence of error in measurements of traffic related air pollution and of blood metabolomics. This kind of formulation allows to model several structures of dependence in a very flexible way, as well as to include an additional component for measurement error. In our work, we applied such methodology to the study of how TRAP measurements are associated with high-throughput molecular data, namely metabolic features sampled from the exposed individuals in a randomized crossover trial. Our application to the Oxford Street II study showed that the inclusion of a classical error term in the models resulted in corrections of the regression estimates whose extent and direction was not evident a priori, thus underlining the importance of explicitly modeling the error component rather than predicting its effect based on prior beliefs. Regression estimates were also corrected by the inclusion of dependency structures in the models, namely dependency among different omic signals and among pollutants. The explicit formulation of such models was possible thanks to the flexible structure of Bayesian hierarchical models, and it was relatively straightforward to embed dependency and measurement error correction in the same hierarchical structure.

This is certainly a major advantage of using Bayesian hierarchical models, which provide a general adaptable way to formulate a broad range of models and structures, as in our case measurement error or dependency structures, and more generally any additional random effect which might need modeling in the analysis. Moreover, the use of a Bayesian framework allows to incorporate prior knowledge in the analysis, for example about the error component and parameters, as well as to reflect the prior uncertainty in the posterior distributions of parameters of interest. This requires some knowledge about the error component, in order to properly formulate the measurement error level and to assign reasonable priors. Note that this requirement is not specific to the Bayesian framework, but rather to any error modeling strategies. In fact, it is always necessary to know the error structure (i.e. classical measurement error and its distribution), as well as the error variance, in order to formulate an identifiable error model (Gustafson 2005). In practice, it is not always straightforward to obtain such information, and often assumptions about the error distribution and parameters are vague or potentially incorrect. The advantage of Bayesian models is that the uncertainty in such assumptions can easily be accounted for and propagated to posterior distributions of corrected estimates.

In comparison to general traditional analyses of the association between metabolite levels and air pollution, and specifically to the work of van Veldhoven et al. (2019), our models allow to account for the presence of measurement error. As a consequence, the estimates of associations between metabolite levels and TRAP exposure are corrected and their absolute values are substantially higher., i.e. they indicate a stronger effect of TRAP on metabolomic features compared with no error adjustment. Moreover, -instead of considering each omic signal separately, we propose to model different signals jointly, so that their correlation is accounted for explicitly. The inclusion of a correlation among exposures is also new with respect to van Veldhoven et al. (2019). While traditional analyses are useful to select candidate signals associated with TRAP exposures, the models we suggest in this work explore the texture of this association in more detail, and correct for biases and correlation structures that might distort their quantification and direction. The models we propose use a Bayesian framework to model measurement error and correlation structures based on the data at hand and to incorporate prior beliefs with the evidence given by the observed data. Doing so, it is possible to account for dependencies among different omics and among pollutants, which can sometimes obscure associations between them and which were not taken into consideration in the traditional model, where associations were assessed separately for each omic signal with each pollutant. Moreover, we included a term for measurement error, which was not accounted for in the traditional model and which can also lead to a biased estimation of these associations.

We implemented our models using `JAGS`, as well as `INLA`. In terms of implementation and computation time, `INLA` performed faster than traditional `MCMC` samplers, specifically `JAGS`, but it showed more issues in the matter of space and memory burdens. In particular, when considering dependency among different omic signals, it was possible to implement the model in `INLA` with only up to 4 signals, while a substantially higher number of signals were included in the `JAGS` model.

With such framework, it is also possible to treat more general models, for example in terms of error structures. It is indeed sufficient to modify the error level in the hierarchical structure, for instance into a Berkson model. Moreover, given the general formulation of Bayesian hierarchical models, it is possible to include more complex dependency structures, for example spatial correlation between pollutants, which can be easily included in the Bayesian structure, especially with `INLA` (Blangiardo et al. 2013). Another possible extension of the method can also be the inclusion of different types of molecular data, which would require a specific and more complex dependency structure among omic signals.

## Acknowledgments

This work was supported by the grant FP7 of the European Commission ‘Enhanced exposure assessment and omic profiling for high priority environmental exposures in Europe’ (EXPOsOMICS grant 308610 to PV). Erica Ponzi was funded by a Doc.Mobility grant from the Swiss National Science Foundation (SNSF grant P1ZHP2_178207).

**Contributions**
E.P. and M.B. conceived the research idea. E.P. designed and conducted the analyses. P.V. and K. F. C. collected and compiled the Oxford Street II data. E.P. and M. B. wrote the manuscript, with inputs from P.V. All authors gave final approval.

## References

Alexeeff, S.E., R.J. Carroll, and B. Coull. 2016. “Spatial Measurement Error and Correction by Spatial SIMEX in Linear Regression Models When Using Predicted Air Pollution Exposures.” Biostatistics 17: 377–89.

Baker, D., and M Nieuwenhuijsen. 2008. Environmental Epidemiology: Study Methods and Applications. Edited by Mark Nieuwenhujisen DeanBaker. DeanBaker, MarkNieuwenhuijsen.

Baxter, L. K., R. J. Wright, C. J. Paciorek, F. Laden, H. H. Suh, and J. I. Levy. 2010. “Effects of Exposure Measurement Error in the Analysis of Health Effects from Traffic-Related Air Pollution.” Journal of Exposure Science and Environmental Epidemiology 20: 101–11.

Blangiardo, M., M. Cameletti, G. Baio, and H. Rue. 2013. “Spatial and Spatio-Temporal Models with R-INLA.” Spatial and Spatio-Temporal Epidemiology 7: 39–55.

Carroll, R.J., D. Ruppert, L.A. Stefanski, and C.M. Crainiceanu. 2006. Measurement Error in Nonlinear Models, a Modern Perspective. Boca Raton: Chapman; Hall.

Coker, E., S. Liverani, J. K. Ghosh, M. Jerrett, B. Beckermann, A. Li, B. Ritz, and J. Molitor. 2016. “Multi-Pollutant Exposure Profiles Associated with Term Low Birth Weight in Los Angeles County.” Environment International 91: 1–13.

Edwards, J.K., and A.P. Keil. 2017. “Measurement Error and Environmental Epidemiology: A Policy Perspective.” Current Environmental Health Reports 4 (1): 79–88.

Elliott, P. J., Cuzick, D. English, and R. Stern. 1992. Geographical and Environmental Epidemiology: Methods for Small-Area Studies. Edited by Oxford University Press. New York, NY: Oxford Univ Press.

Goldman, G.T., J.A. Mulholland, A. G. Russell, M.J. Strickland, M. Klein, L.A. Waller, and P.E. Tolbert. 2011. “Impact of Exposure Measurement Error in Air Pollution Epidemiology: Effect of Error Type in Time-Series Studies.” Environmental Health 10 (61).

Gryparis, A., B.A. Coull, and J. Schwarzt. 2007. “Controlling for Confounders in the Presence of Measurement Error in Hierarchical Models; a Bayesian Approach.” Journal of Exposure Science and Environmental Epidemiology 17: S20–S28.

Gryparis, A., A. Zeka, J. Schwartz, and B.A. Coull. 2009. “Measurement Error Caused by Spatial Misalignment in Environmental Epidemiology.” Biostatistics 10: 258–74.

Gustafson, P. 2005. “On Model Expansion, Model Contraction, Identifiability and Prior Information: Two Illustrative Scenarios Involving Mismeasured Variables.” Statistical Science 20: 111–40.

Huang, G., and E. M. Lee D. amd Scott. 2018. “Multivariate Space-Time Modelling of Multiple Air Pollutants and Their Health Effects Accounting for Exposure Uncertainty.” Statistics in Medicine 37: 1134–48.

Mallick, B., F.O. Hoffman, and R.J. Carroll. 2002. “Semiparametric Regression Modeling with Mixtures of Berkson and Classical Error, with Application to Fallout from the Nevada Test Site.” Biometrics 58: 13–20.

Newton, M.A., A. Noueiry, D. Sarkar, and P. Ahlquist. 2004. “Detecting Differential Gene Expression with a Semiparametric Hierarchical Mixture Method.” Biostatistics 5 (2): 155–76.

Pekkanen, J., and N. Pearce. 2001. “Environmental Epidemiology: Challenges and Opportunities.” Environmental Health Perspectives 109: 1–5.

Rhomberg, L.R., J.K. Chandalia, C. Long, and J.E. Goodman. 2011. “Measurement Error in Environmental Epidemiology and the Shape of Exposure-Response Curves.” Critical Reviews in Toxicology 41 (8): 651–71.

Richardson, S., and W.R. Gilks. 1993. “Conditional Independence Models for Epidemiological Studies with Covariate Measurement Error.” Statistics in Medicine 12: 1703–22.

Rue, H, S Martino, and N Chopin. 2009. “Approximate Bayesian Inference for Latent Gaussian Models by Using Integrated Nested Laplace Approximations (with Discussion).” Journal of the Royal Statistical Society Series B (Statistical Methodology) 71: 319–92.

Sinharay, R., J. Gong, B. Barratt, P. Ohman-Strickland, S. Ernst, F. Kelly, J.J. Zhang, P. Collins, P. Cullinan, and K.F. Chung. 2017. “Respiratory and Cardiovascular Responses to Walking down a Traffic-Polluted Road Compared with Walking in a Traffic-Free Area in Participants Aged 60 Years and Older with Chronic Lung or Heart Disease and Age-Matched Healthy Controls: A Randomised,Crossover Study.” Lancet 391: 339–49.

Stephens, D.A., and P. Dellaportas. 1992. “Bayesian Analysis of Generalised Linear Models with Covariate Measurement Error.” In Bayesian Statistics 4, edited by J. M. Bernardo, J. O. Berger, A. P. Dawid, and A. F. M. Smith. Oxford Univ Press.

Van Roosbroeck, S., R. Li, G. Hoek, E. Lebret, B. Brunekeef, and D. Spiegelmann. 2006. “Traffic-Related Outdoor Air Pollution and Respiratory Suymptoms in Children: The Impact of Adjustment for Exposure Measurement Error.” Epidemiology 368: 174–84.

Veldhoven, K. van, P. Keksi-Rahkonen, A. Kiss, A. Scalbert, P. Cullinan, F. Chung, B. Barrat, et al. 2019. “Impact of Short-Term Traffic-Related Air Pollution on the Metabolome-Results from Two Metabolome-Wide Experimental Studies.”, Environment International 123: 124–131.

Ventrucci, M., E.M. Scott, and D. Cocchi. 2011. “Multiple Testing on Standard Mortality Ratios: A Bayesian Hierarchical Model for FDR Estimation.” Biostatistics 12 (1): 51–67.

Vineis, P, M Chadeau-Hyam, H Gmuender, J Gulliver, Z Herceg, J Kleinjans, M Kogevinas, et al. 2017. “The Exposome in Practice: Design of the EXPOsOMICS Project.” International Journal of Hygiene and Environmental Health 220: 142–51.

Zeger, S.L., D. Thomas, F. Dominici, J.M. Samet, J. Schwartz, D. Dockery, and A. Cohen. 2000. “Exposure Measurement Error in Time-Series Studies of Air Pollution: Concepts and Consequences.” Environmental Health Perspectives 108: 419–26.

